# Bayesian inference and comparison of stochastic transcription elongation models

**DOI:** 10.1101/499277

**Authors:** Jordan Douglas, Richard Kingston, Alexei J. Drummond

## Abstract

Transcription elongation can be modelled as a three step process, involving polymerase translocation, NTP binding, and nucleotide incorporation into the nascent mRNA. This cycle of events can be simulated at the single-molecule level as a continuous-time Markov process using parameters derived from single-molecule experiments. Previously developed models differ in the way they are parameterised, and in their incorporation of partial equilibrium approximations.

We have formulated a hierarchical network comprised of 12 sequence-dependent transcription elongation models. The simplest model has two parameters and assumes that both translocation and NTP binding can be modelled as equilibrium processes. The most complex model has six parameters makes no partial equilibrium assumptions. We systematically compared the ability of these models to explain published force-velocity data, using approximate Bayesian computation. This analysis was performed using data for the RNA polymerase complexes of *E. coli, S. cerevisiae* and Bacteriophage T7.

Our analysis indicates that the polymerases differ significantly in their translocation rates, with the rates in T7 pol being fast compared to *E. coli* RNAP and *S. cerevisiae* pol II. Different models are applicable in different cases. We also show that all three RNA polymerases have an energetic preference for the posttranslocated state over the pretranslocated state. A Bayesian inference and model selection framework, like the one presented in this publication, should be routinely applicable to the interrogation of single-molecule datasets.

**Author summary:** Transcription is a critical biological process which occurs in all living organisms. It involves copying the organism’s genetic material into messenger RNA (mRNA) which directs protein synthesis on the ribosome. Transcription is performed by RNA polymerases which have been extensively studied using both ensemble and single-molecule techniques (see reviews: [1, 2]). Single-molecule data provides unique insights into the molecular behaviour of RNA polymerases. Transcription at the single-molecule level can be computationally simulated as a continuous-time Markov process and the model outputs compared with experimental data. In this study we use Bayesian techniques to perform a systematic comparison of 12 stochastic models of transcriptional elongation. We demonstrate how equilibrium approximations can strengthen or weaken the model, and show how Bayesian techniques can identify necessary or unnecessary model parameters. We describe a framework to a) simulate, b) perform inference on, and c) compare models of transcription elongation.

## Introduction

Transcription is carried out by RNA polymerases: RNAP in *Escherichia coli*, pol II in *Saccharomyces cerevisiae*, and T7 pol in Bacteriophage T7. It involves the copying of template double-stranded DNA (dsDNA) into single-stranded messenger RNA (mRNA). RNAP and pol II are comprised of multiple subunits, and their catalytic subunits are homologous [3,4]. In contrast, T7 pol exists as a monomer with a distinct sequence, and resembles the *E. coli* DNA polymerase I [5].

Optical trapping experiments have been performed on the transcription elongation complex (TEC) from a variety of organisms [6–12]. In a typical experimental setup, two polystyrene beads (around 600 nm in diameter) are tethered to the system; one attached to the RNA polymerase and the other to the DNA [6]. As transcription elongation progresses, the distance between the two beads increases and the velocity of a single TEC can be computed. Optical tweezers can be used to apply a force *F* to the system (Fig 1).

**Fig 1.**
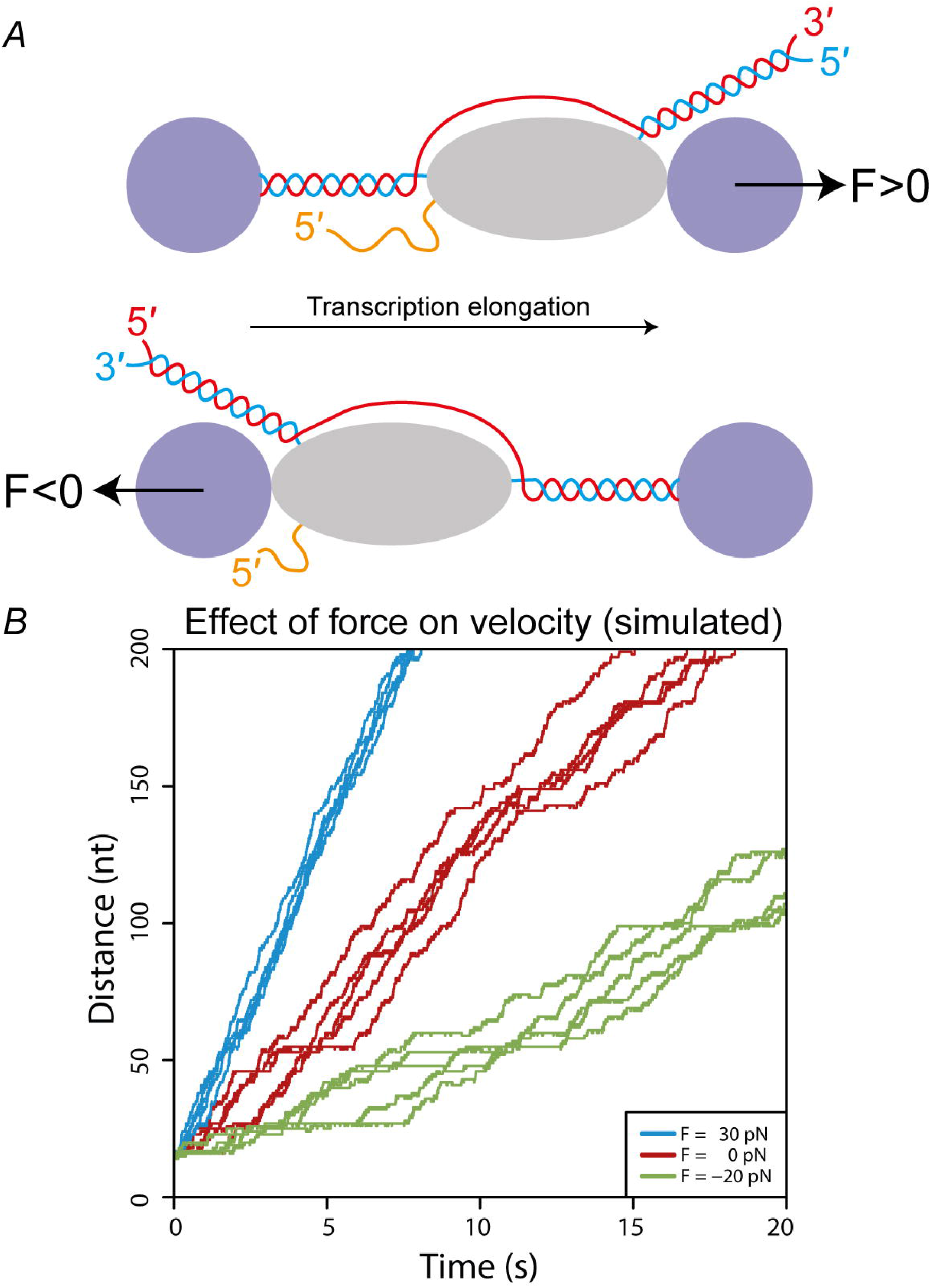
Effect of an applied force on elongation velocity. (A) Optical trapping setup showing dsDNA being transcribed by RNA polymerase (grey ellipse) into mRNA. Two polystyrene beads are tethered to the system allowing the application of force using optical tweezers. An assisting load *F* > 0 acts in the same direction as transcription (top) while a hindering load *F* < 0 acts in the opposing direction (bottom). Figure not to scale. (B) Schematic depiction of the effect of applying a force on RNA polymerase. Due to the stochastic nature of transcription at the single-molecule level, each experiment yields a different distance-time trajectory, even under the same applied force.

Single-molecule studies of the TEC have revealed that RNA polymerases progress in a discontinuous fashion [6,13–16] with step sizes that correspond to the dimensions of a single nucleotide (3.4 Å [17]). Consequently, at the single molecule level, transcription is best modelled as a discrete process rather than a continuous one.

A single cycle in the main transcription elongation pathway (Fig 2) requires (1) Forward translocation of the RNA polymerase, making the active site accessible; (2) Binding of the complementary nucleoside triphosphate (NTP); (3) Addition of the nucleotide onto the 3′ end of the mRNA. This third step involves NTP hydrolysis. Nucleoside monophosphate is added onto the chain and pyrophosphate is released from the enzyme.

**Fig 2.**
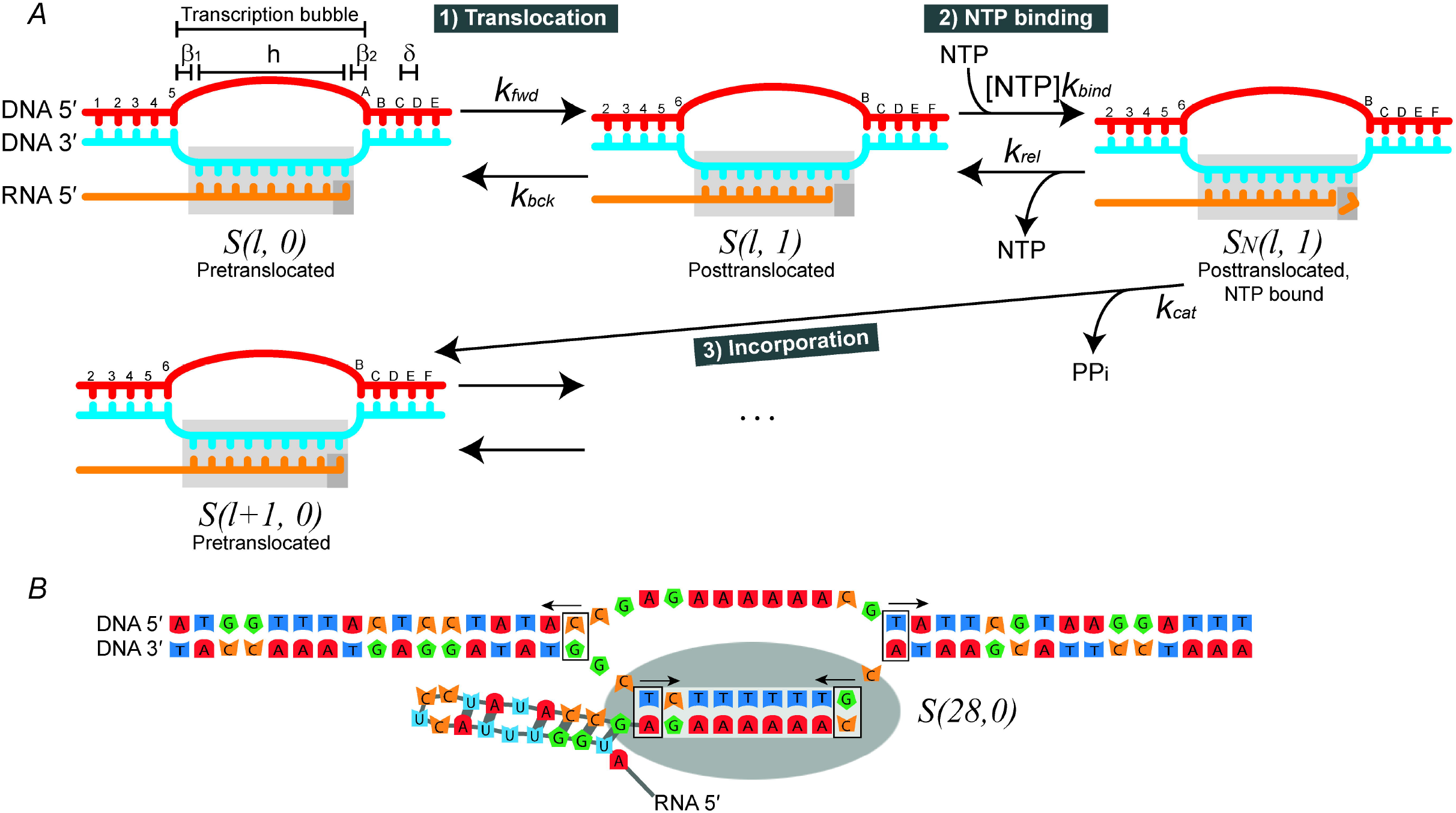
State diagrams of RNA polymerase. (A) The model of the main transcription elongation pathway, which shows the postulated states; the pathways for interconversion; and the rate constants that govern each part of the reaction. The transcription bubble is the set of *β*_1_ + *h* + *β*_2_ bases (see main text for definitions) in the double-stranded DNA which are unpaired. States are denoted by *S*(*l,t*) where *l* is the length of the mRNA and *t* is the position of the polymerase active site (small grey rectangle) with respect to the 3′ end of the mRNA. Polymerase translocation displaces the polymerase by a distance of *δ* = 1 bp = 3.4 Å. During polymerisation the chain is extended by one nucleotide. (B) Instantiated posttranslocated state of RNA polymerase transcribing the *rpoB* gene sequence, with *β*_1_ = 2, *h* = 9, *β*_2_ =1. Forward translocation requires melting two T/A basepairs (right arrows). Backward translocation requires melting two C/G basepairs (left arrows). The mRNA secondary structure would also require reconfiguration [18,19].

Our study aimed to identify the best model to describe this reaction cycle for RNAP, pol II, and T7 pol, based on analysis of published force-velocity data. As there are three reactions, up to six rate constants may be necessary for a kinetic model of a single nucleotide addition. These describe forward and backwards translocation (*k_fwd_* and *k_bck_*), binding and release of NTP (*k_bind_* and *k_rel_*), and NTP catalysis and reverse-catalysis (*k_cat_* and *k_rev_*), also known as pyrophosphorolysis [20]. However fewer than six parameters may be required in practice.

First, it is reasonable to assume that polymerisation is effectively irreversible [19,21–23], as pyrophosphorolysis is a highly exergonic reaction, reducing the number of rate constants to five. Second, translocation between the pretranslocated and posttranslocated states, and/or NTP binding, may occur on timescales significantly more rapid than the other steps, in which case they may be modelled as equilibrium processes. These assumptions simplify the model, as the respective forward and reverse reaction rate constants are subsumed by a single equilibrium constant. Third, thermodynamic models of nucleic acid structure can be used to estimate sequence-dependent translocation rates *k_fwd_*(*l*) and *k_bck_*(*l*), by invoking transition state theory, and this can sometimes result in parameter reduction [18,19,23].

Irrespective of equilibrium assumptions and parameterisation, transcription elongation under applied force can be modelled in two fundamentally distinct ways. First, there are the **deterministic** equations which can be used to calculate the mean pause-free elongation velocity *v*(*F*, [NTP]) as a function of force *F* and NTP concentration [NTP]. This kind of model can be derived from the differential equations describing the time evolution of all species, by application of the steady state approximation. Force effects on the translocation step are incorporated using transition state theory [24,25].

An example is the following 3-parameter model [6].

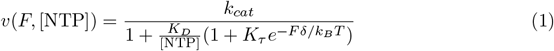

where *δ* is the distance between adjacent basepairs (3.4 Å, [17]), 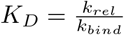 is the equilibrium constant of NTP binding, 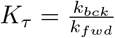 is the equilibrium constant of translocation, *k_B_* is the Boltzmann constant, and *T* is the absolute temperature. Increasingly complex equations may be used as more parameters or states are added to the model [6, 8, 19]. Such equations describe the velocity averaged across an ensemble of molecules. Parameter inference applied to velocity-force-[NTP] experimental data is straightforward and computationally fast when using these equations. However these equations do not describe the distribution of velocity nor do they account for site heterogeneity across the nucleic acid sequence and therefore cannot predict local sequence effects.

Second, there are the **stochastic** models, which can be implemented via simulation of single-molecule behaviour using the Gillespie algorithm [26]. The mean velocity can be calculated by averaging velocities over a number of simulations for a given *F* and [NTP]. This offers not just the mean but a full distribution of velocities and could potentially explain emergent properties unavailable from a deterministic model. Unfortunately, simulating can be very slow and therefore parameter inference can be a problem.

In this study we used a Markov-chain-Monte-Carlo approximate-Bayesian-computation (MCMC-ABC) algorithm [27] to estimate transcription elongation parameters for stochastic models via simulation. The observed pause-free velocities we are fitting to were measured at varying applied force and NTP concentration. For each RNA polymerase under study - *E. coli* RNAP, *S. cerevisiae* pol II, and T7 pol - we fit to one respective dataset from the single-molecule literature [6,28,29].

## Models

### Notation and state space

Suppose the TEC is transcribing a gene of length *L*. Then let *S*(*l, t*) denote a TEC state, where the mRNA is currently of length *l* ≤ *L*, and 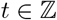 describes the position of the active site with respect to the 3′ end of the mRNA. When *t* = 0 the polymerase is pretranslocated and cannot bind NTP, and when *t* = 1 the polymerase is posttranslocated and *can* bind NTP (Fig 2). This study is focused on the main elongation pathway and the observed velocities being fitted have pauses filtered out. Therefore, although additional backtracked states (*t* < 0) [6,30,31] and hypertranslocated states (*t* > 1) [32,33] exist, these are not incorporated in the model.

Let *β*_1_ and *β*_2_ be the number of unpaired template nucleotides upstream and downstream of RNA polymerase, respectively, and let *h* be the number of basepairs in the DNA/mRNA hybrid (Fig 2A). Although there are uncertainties in these parameters, they are held constant at *h* = 9, *β*_1_ = 2, and *β*_2_ = 1 [19,34].

Transcription of the gene begins at state *S*(*l*_0_, 0) and ends upon reaching *S*(*L*, 0), where *l*_0_ = *β*_1_ + *h* + 2.

### Parameterisation of the NTP binding step

NTP binding has been modelled as both a kinetic and equilibrium process in the literature [6,19,23].

In a kinetic binding model, NTP binding occurs at pseudo-first order rate *k_bind_*[NTP], while NTP release occurs at rate *k_rel_*. In this case, *k_bind_* and 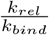 must be estimated.

Under a partial equilibrium approximation NTP binding and release are assumed to be rapid enough that equilibrium is achieved. In this case, the rate constants *k_bind_* and *k_rel_* are subsumed by the NTP dissociation constant 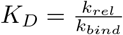 which becomes the sole binding-related parameter to estimate.

### Parameterisation of the translocation step

While inferences about the rate constants associated with NTP binding and catalysis (*k_bind_*, 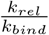, and *k_cat_*) can be made directly from the data, the translocation step is more complex. Transition state theory is invoked in order to estimate *k_fwd_* and *k_bck_*. Recasting the problem in this way (1) provides a way of accommodating the effects of applied force on the elongation process, and (2) allows the sequence-dependence of translocation to be incorporated by considering the energetics of basepairing. When allowing for sequence dependence, the total number of translocation rates required to model translocation of the full gene is 2(*L* − *l*_0_).

### Thermodynamic models of base pairing energies

The standard Gibbs free energies Δ_*r*_*G*^0^(= Δ*G*) involved in duplex formation are calculated using nearest neighbour models. The standard Gibbs energy of state *S* – arising from nucleotide basepairing and dangling ends – is calculated as

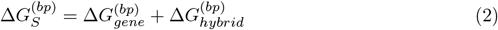

where SantaLucia’s DNA/DNA basepairing parameters [35] are used to calculate 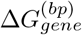 and Sugimoto’s DNA/RNA parameters [36] are used for 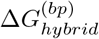. For the latter, dangling end energies are estimated as described by Bai et al. 2004 [23]. Here, and elsewhere, the^(*bp*)^ superscript is used to denote a model parameter that can be evaluated from the sequence alone. Gibbs energies are expressed relative to the thermal energy of the system, in units of *k_B_T*, where *k_B_T* = 0.6156 kcal/mol at *T* = 310 K.

In order for RNA polymerase to translocate forward (backward), up to two basepairs must be disrupted: (1) the basepair at the downstream (upstream) edge of the transcription bubble, and (2) the basepair at the upstream (downstream) end of the DNA/mRNA hybrid (Fig 2B). Differences in the basepairing energies in these regions confer sequence-dependence on the rate of translocation.

### Calculation of translocation rates or translocation equilibrium constant

The standard Gibbs energies of the pre and posttranslocated states, 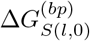 and 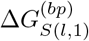, respectively, are used with up to four additional terms – Δ*G*_*τ*1_, *δ*_1_, 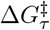, and 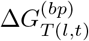 – to calculate the translocation rates. The first three are model parameters which must be estimated while the latter is directly evaluated from the sequence.

Let *T*(*l,t*) be the translocation transition state between *S*(*l,t*) and *S*(*l,t* + 1). Then 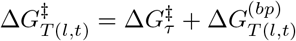 is the sequence-dependent standard Gibbs energy of activation which must be overcome in order to translocate (Fig 3).

**Fig 3.**
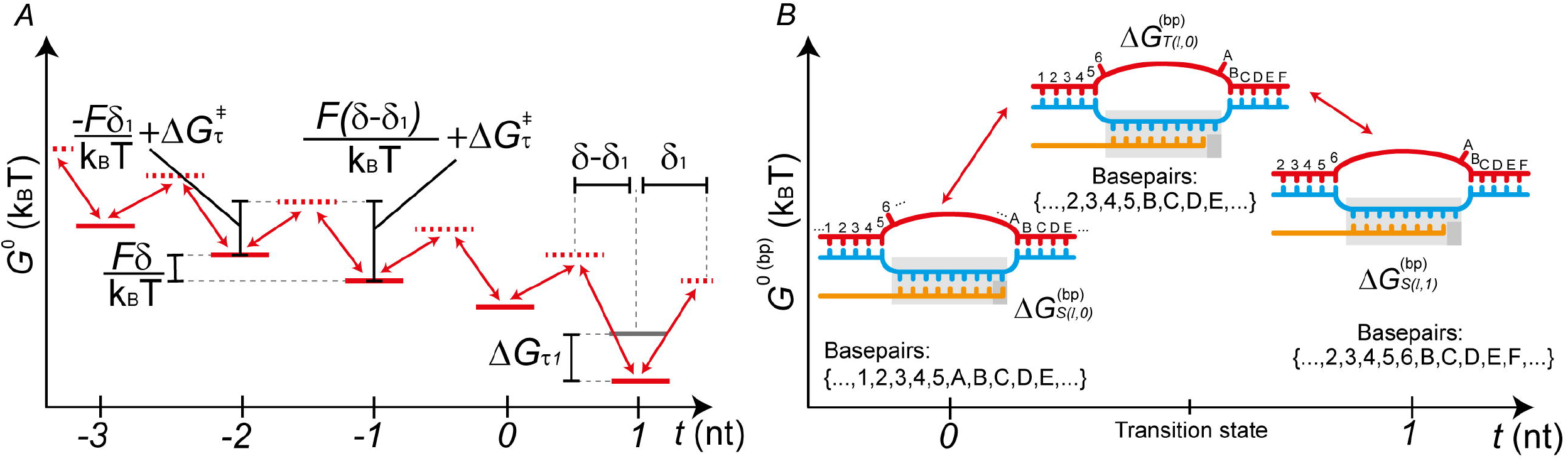
Parameterisation of the translocation step. (A) Effects of model parameters on state energies. The figure displays a schematic Gibbs energy landscape of translocation, with backtracked states included for visualisation purposes. The solid red lines represent translocation states (*t* = 0: pretranslocated, *t* =1: posttranslocated, and *t* < 0: backtracked), while the dashed red lines represent transition states. Applying an assisting force *F* > 0 tilts the landscape in favour of higher values of *t*. The effect of Δ*G*_*τ*1_ is observed at the posttranslocated state *t* = 1. In a translocation equilibrium model, the barrier height is assumed to be so small, = translocation is so rapid, that the transition states are disregarded. (B) A model for the sequence-dependent transition state between translocation states *S*(*l*, 0) and *S*(*l*, 1). This is required for estimating the Gibbs energy of basepairing 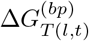 in the transition state. The basepairing energy, added to a baseline term 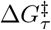, together specify the height of the activation barrier (Equation 10).

Given an applied force *F*, the translocation rates governing transition between the pre and posttranslocated states (*k_fwd_*(*l*) and *k_bck_*(*l*)) are calculated from barrier height 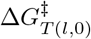 using an Arrhenius type relation:

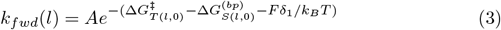

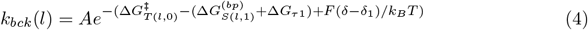

The derived rates *k_fwd_*(*l*) and *k_bck_*(*l*) are therefore dependent on the local sequence. The pre-exponential factor *A* is held constant at 10^6^ s^−1^. This term has been arbitrarily set to a variety of values in previous studies (10^6^ – 10^9^ s^−1^ [18,19,23]). This has little consequence for model fitting, however the value of 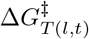 is entangled with the value of the pre-exponential factor *A* and can only be meaningfully interpreted in light of its value.

If the system has time to reach equilibrium, the probabilities of observing the pretranslocated state *S*(*l*, 0) and posttranslocated state *S*(*l*, 1) are

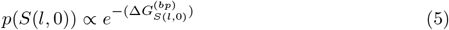

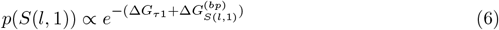

This is described by equilibrium constant *K_τ_*.

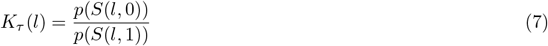

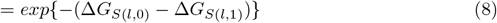

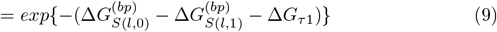

The physical meanings of the terms Δ*G*_*τ*1_, *δ*_1_, 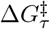, and 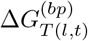, and the way they are used in the model, are detailed below.

### Energetic bias for the posttranslocated states

Δ*G*_*τ*1_ (units *k_B_T*) is a parameter added to the standard Gibbs energy of the posttranslocated state. If Δ*G*_*τ*1_ = 0, then the sequence alone determines the Gibbs energy difference between pre and posttranslocated states. In this case, pretranslocated states are usually favoured over posttranslocated states due to the loss of a single basepair in the hybrid of the latter.

Δ*G*_*τ*1_ has frequently been estimated for T7 pol [37–39] and there has been discussion around whether such a term is necessary for RNAP [8].

### Polymerase displacement and formation of the transition state

*δ*_1_ (units Å) is the distance that the polymerase must translocate forward to facilitate the formation of the transition state. The distance between adjacent basepairs is held constant at an experimentally measured value *δ* = 3.4 Å [17], and 0 < *δ*_1_ < *δ*. The response of the system to an applied force *F* depends on this term. In general, the application of force *F* tilts the Gibbs energy landscape – the Gibbs energy difference between adjacent translocation states being augmented by a factor 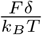 (Fig 3A, [1,40]).

It may be necessary to estimate *δ*_1_ to model the data adequately [19], or it may be sufficient to simply set *δ*_1_ = *δ*/2 [40].

### Energy barrier of translocation

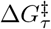 and 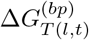 (units *k_B_T*) together determine the activation barrier height in the translocation step. It is assumed that the sequence-dependent standard Gibbs energy of activation 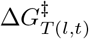 can be written as

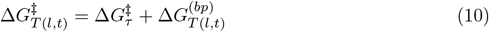

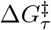 is therefore a sequence-independent baseline term used to compute the translocation barrier heights. The parameter 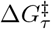 must be estimated in order to evaluate translocation rates.

In contrast 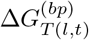 is a term that is evaluated directly from the sequence derived from a model of the transition state (Fig 3B). The term is evaluated as the standard Gibbs energy of a TEC containing all hybrid and gene basepairs found in both *S*(*l, t*) and *S*(*l,t* + 1), ie. the intersection between the two sets of basepairs.

### Model space

The full transcription elongation model makes use of the following 6 parameters:

- *k_cat_* (units s^−1^).
- 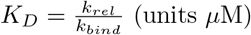.
- *k_bind_* (units *μ*M^−1^ s^−1^).
- Δ*G*_*τ*1_ (units *k_B_T*).
- *δ*_1_ (units Å).
- 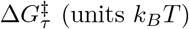.

However fewer than 6 parameters may be needed to adequately describe the data. If it is assumed that the energy differences between pre and posttranslocated states are determined by basepairing energies alone, the parameter Δ*G*_*τ*1_ does not need to be estimated. This is equivalent to holding Δ*G*_*τ*1_ constant at 0. If it is assumed that the displacement required for formation of the translocation intermediate state is half the distance between adjacent basepairs, the parameter *δ*_1_ does not need to be estimated. This is equivalent to holding *δ*_1_ constant at *δ*/2.

Partial equilibrium approximations may also simplify the model, as detailed above. If binding is approximated as an equilibrium process, *k_bind_* does not need to be estimated. If translocation is approximated as an equilibrium process, 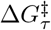 and *δ*_1_ do not need to be estimated. One, both, or neither of these two steps (binding and translocation) could be assumed to achieve equilibrium, thus yielding four equilibrium model variants (Fig 4A). The introduction of partial equilibrium approximations for both the NTP binding and translocation steps has implications when specifying the prior distributions for the Bayesian analysis (S4 Appendix.) The chemical master equations for single nucleotide addition cycles of these models are presented in S2 Appendix.

**Fig 4.**
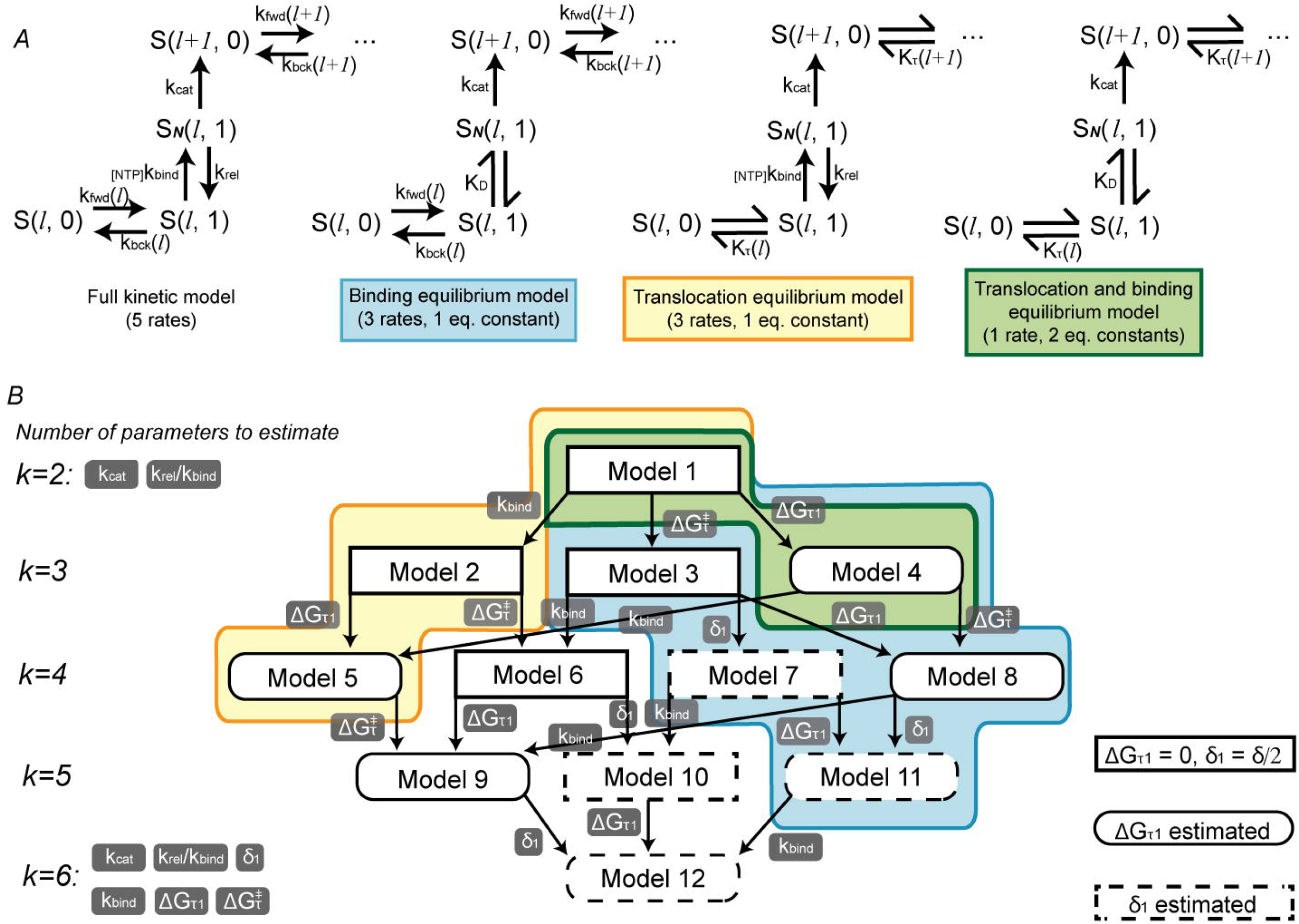
The space of models to be compared. (A) The four equilibrium model variants. NTP binding, translocation, both, or neither, could be assumed to achieve equilibrium prior to catalysis. (B) The 12 transcription elongation models. An arrow connects model *i* to *j* if augmentation of model *i* with a single parameter generates model *j*. The number of parameters to estimate *k* is shown for each level in the network. Equilibrium approximation colour scheme is the same as in A. Δ*G*_*τ*1_ and *δ*_1_ can each be estimated or set to a constant.

Incorporating these simplifications to the model in a combinatorial fashion results in a total of 12 related models, which together constitute the model space. Our objective was to determine which of these 12 models provides the best description of the experimental data. The simplest model (Model 1) contains 2 parameters (*k_cat_* and *K_D_*). The most complex model (Model 12) contains all 6 parameters. The full model space is displayed in Fig 4B.

### Stochastic modelling

For each model we performed stochastic simulations, appropriate for the modelling of single-molecule force-velocity data. The simulations, performed using the Gillespie algorithm [26,41], can be used to estimate the mean elongation velocity under a model.

The estimation of mean velocity can be broken down into three steps. First, the system is initialised by placing the RNA polymerase at the 3′ end of the template – state *S*(*l*_0_, 0) – with the transcription bubble open and a DNA/RNA hybrid formed. The force and NTP concentrations are assigned their experimentally set values. Second, a chemical reaction is randomly sampled. The probability that reaction 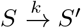 is selected is proportional to its rate constant *k* (Fig 2). The amount of time taken for the reaction to occur is sampled from the exponential distribution. States which are subject to a partial equilibrium approximation are coalesced into a single state, which augments the outbound rate constants. The second step is repeated until the RNA polymerase has copied the entire template. Third, the previous two steps are repeated c times. The mean elongation velocity is evaluated as the mean of each mean elongation velocity across c simulations. For further information, see S1 Appendix.

### Relation to previous models and stochastic simulations

There is an extensive literature concerned with the kinetic modelling of transcription elongation. Such models may incorporate backtracking, hypertranslocation, and other reactions. Here we are concerned only with the central elongation pathway.

A stochastic and sequence-dependent model was proposed by Bai et al. 2004 [23] for RNAP, with both NTP binding and translocation treated as equilibrium processes. The translocation equilibrium constant was calculated entirely from basepairing energies. Therefore this model is equivalent to Model 1, and the parameters were estimated as *k_cat_* = 24.7 s^−1^ and *K_D_* = 15.6 *μM* from fit to experimental data. Maoiléidigh et al. 2011 also presented stochastic simulations of RNAP. The elongation component of their model is equivalent to Model 6 [19]. We build on this work by providing a systematic Bayesian framework for model comparison and parameter estimation.

While our analysis employed sequence-dependent stochastic models, comparisons can also be made with some deterministic models.

Abbondanzieri et al. 2005 [6], Larson et al. 2012 [42], Schweikhard et al. 2014 [28], and Thomen et al. 2008. [29,39] described a deterministic model (for RNAP, pol II, pol II, and T7 pol respectively) which estimated *k_cat_, K_D_* and translocation equilibrium constant 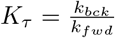. These are most similar to Model 4.

Maoiléidigh et al. 2011 for RNAP, and Dangkulwanich et al. 2013 for pol II, however found that the translocation and catalysis were occurring on similar timescales, and modelled only NTP binding as an equilibrium process [19, 43]. They also estimated the distance of translocation. These deterministic models are most similar to Model 11.

Finally, Mejia et al. 2015 [44] used a model that is quite different to all the above models, as it does not explicitly treat translocation. Instead elongation is modelled with a two step kinetic scheme, the first step involving NTP binding and conformational change, and the second step involving nucleotide incorporation and product release. This model is most similar to a special case of Model 5 where Δ*G*_*τ*1_ becomes extremely negative, driving the polymerase into the posttranslocated position.

## Results and Discussion

### Model selection with MCMC-ABC

Our aim was to 1) use Bayesian inference to select the best of 12 transcription elongation models for each RNA polymerase; and 2) estimate the parameters for those of the models appearing in the 95% credible set of the posterior distribution. Selecting prior distributions behind each parameter is a critical process in Bayesian inference. A prior distribution should reflect what is known about the parameter before observing the new data. We have explicated our prior assumptions, with justifications, in Table 1.

**Table 1.** Prior distributions used during Bayesian inference.

| Parameter | Prior distribution(s) | Justification of prior distribution(s) |
| --- | --- | --- |
| Model $M$ | $P(M_i) = 2/16$ for $i \in \{1, 2, 4, 5\}$<br>$P(M_i) = 1/16$ for $i \in \{3, 6, 7, 8, 9, 10, 11, 12\}$ | Each model should each have uniformly distributed values.<br>Models with translocation at equilibrium have double the prior probability since these models do not use $\delta_1$. |
| $k_{cat}$ ($s^{-1}$) | Lognormal($\mu = 3.454$, $\sigma = 0.587$) **for RNAP/pol II**<br>Lognormal($\mu = 4.585$, $\sigma = 0.457$) **for T7 pol** | $k_{cat}$ and elongation velocity estimates for *E. coli* RNAP and *S. cerevisiae* pol II range from 18 to 50 $s^{-1}$ for optical trapping experiments [8–10, 23, 44], but as much as 100 bp/s *in vivo* [46–49]. Distribution selected such that (10, 100) is central 95% interval.<br>For T7 pol $k_{cat}$ and elongation velocity estimates range from 43 - 240 bp/s [11, 50–52]. Distribution selected such that (40, 240) is central 95% interval. |
| $K_D$ ($\mu M$) | Lognormal($\mu = 1.844$, $\sigma = 1.762$) | Estimates for $K_D$ under binding equilibrium models range from 20-140 $\mu M$ [8, 22, 39, 42, 53]. In models where binding is kinetic and slow, $K_D \equiv \frac{k_{rel}}{k_{bind}}$ could be much lower (S4 Appendix). To accommodate for both binding models, the prior distribution was selected such that the central 95% interval is (0.2, 200). |
| $k_{bind}$ ($\mu M^{-1}s^{-1}$) | Lognormal($\mu = -1.498$, $S\sigma = 1.585$) | Central 95% interval set so that NTP binding is a slow kinetic step (S4 Appendix ). Centered around (0.01, 5). |
| $\Delta G_{\tau 1}$ ($k_B T$) | Normal($\mu = 0$, $\sigma = 1.55$) **for RNAP/pol II**<br>Normal($\mu = -3.3$, $\sigma = 1.55$) **for T7 pol** | For RNAP and pol II, centered around 0 with a standard deviation comparable to the free energy of a single nucleotide basepair doublet, and such that the 95% central interval is (-4, 4).<br>For T7 pol $\Delta G_{\tau 1}$ has been estimated as -4.3 [39] and -4.87 $k_B T$ [37]. However these estimates are likely resulting partially from dangling ends. Thus, we subtracted the mean dangling end contribution of $\sim -1$ $k_B T$ [35] and centered the prior around this interval with a standard deviation the same as above. |
| $\Delta G_{\tau}^{\ddagger}$ ($k_B T$) | Normal($\mu = 5.5$, $\sigma = 0.97$) **for RNAP/pol II**<br>Normal($\mu = 2.5$, $\sigma = 1.36$) **for T7 pol** | Central 95% interval set so that translocation is a slow kinetic step (S4 Appendix ). Selected so that 99% central interval is (3, 8) for RNAP and pol II, and (-1, 6) for T7 pol. |
| $\delta_1$ ($\text{\AA}$) | Uniform($l = 0$, $u = 3.4$) | Uniformly distributed across all possible values. |

Prior distributions behind all estimated parameters and the model indicator. Unless specified otherwise, the prior distribution is used for all three RNA polymerases. Lognormal priors (parameterised in log space) are used for rates and equilibrium constants while normal priors are used for Gibbs energy terms. To maintain statistical integrity of the Bayesian analysis, prior distributions were not derived from the data presented by Abbondanzieri et al. 2005 [6] for RNAP, by Schweikhard et al. 2014 [28] for pol II, or by Thomen et al. 2008 [29] for T7 pol.

We performed MCMC-ABC experiments which estimated the parameters and model indicator *M_i_* for 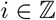, 1 ≥ *i* ≥ 12. Models which appear more often in this posterior distribution are better choices, given the data.

The datasets we fit our models to are all from the single-molecule literature and are presented in: Figures 5a and 5b of Abbondanzieri et al. 2005 [6] for *E. coli* RNAP, Figure 2a of Schweikhard et al. 2014 [28] for *S. cerevisiae* pol II, and Table 2 of Thomen et al. 2008 [29] for T7 pol. To computationally replicate these experiments as faithfully as we could with the available information and computational limitations, simulations in this study were run on the 4 kb *E. coli rpoB* gene for RNAP (GenBank: EU274658), the first 4.75 kb of the human *rpb1* gene for pol II (NCBI: NG_027747) the: first 10 kb of the Enterobacteria phage λ genome for T7 pol (NCBI: NC_001416). The: mean velocities from 32 (for RNAP), 10 (for pol II) and 3 (for T7 pol) simulations of the full respective sequences were used to estimate the mean elongation velocity during MCMC-ABC, given *F* and [NTP].

**Fig 5.**
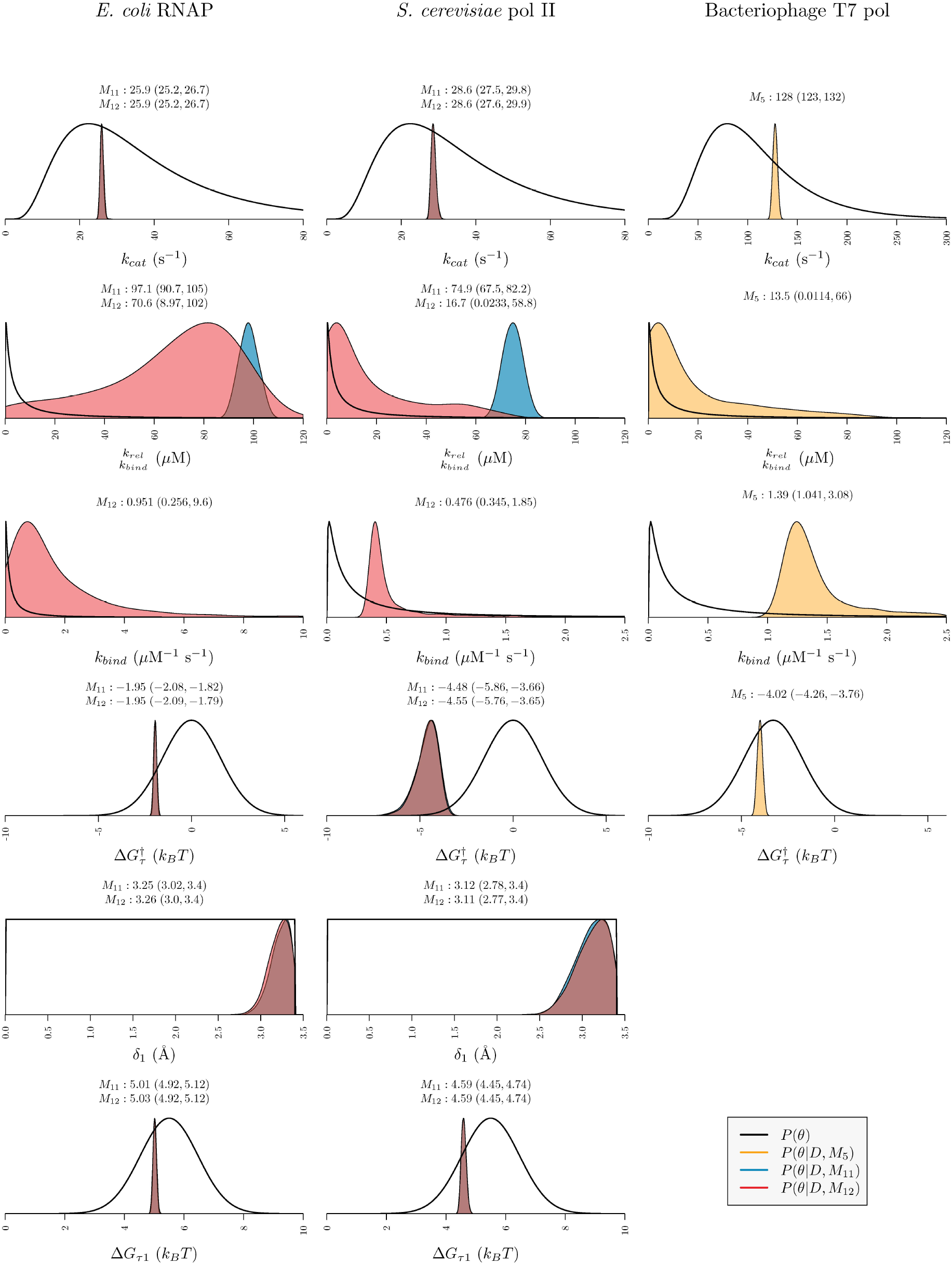
Posterior and prior distribution plots. Posterior distributions for all models which appear in the 95% credible set are displayed (two models for RNAP, two models for pol II, and one model for T7 pol). Plots show the prior probability density *P*(*θ*) of each parameter and posterior probability density of each parameter conditional on the model *P*(*θ*|*D*, *M_i_*). The geometric median point-estimates and highest posterior density (HPD) intervals (calculated with Tracer 1.6 [54]) are displayed above each plot (3 sf).

**Table 2.**
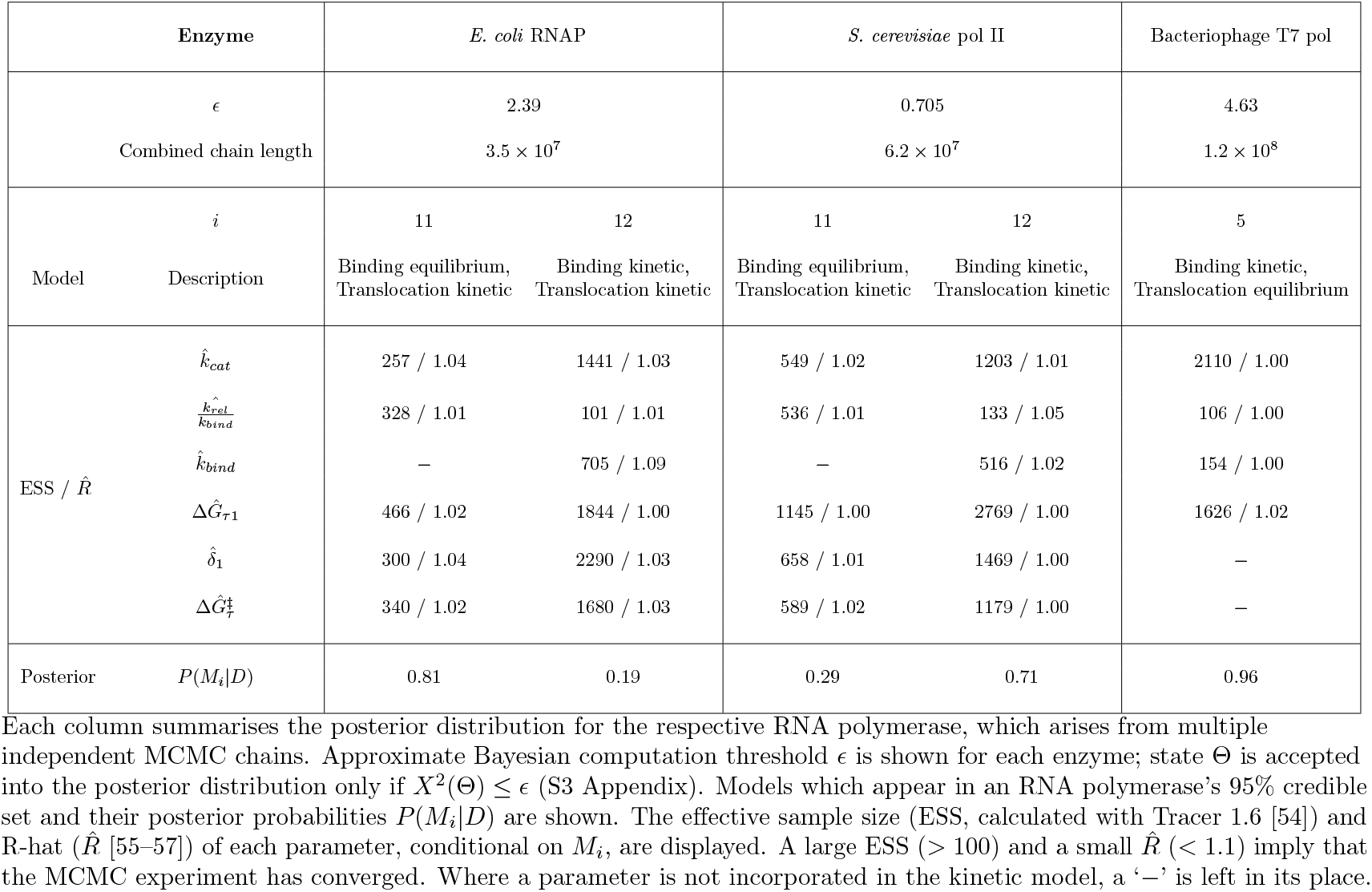
Summary of MCMC-ABC experiments.

For further information about the MCMC-ABC algorithm [27,45], see S3 Appendix.

### The posterior distributions

The posterior distributions from our MCMC-ABC experiments are presented in Table 2, Fig 5, and Fig 6.

**Fig 6.**
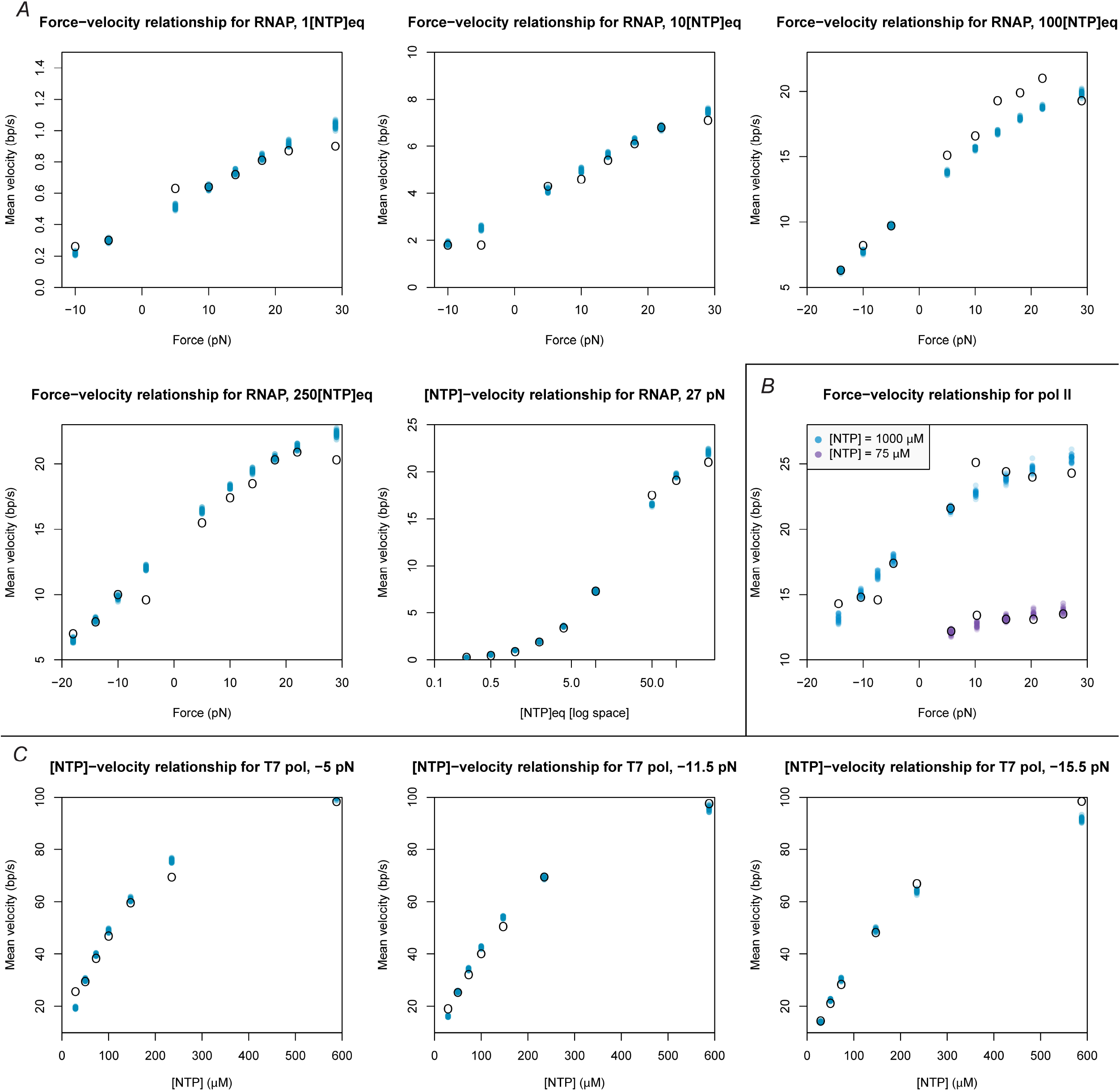
Posterior distributions of simulated velocities. Black open circles represent experimentally measured mean velocities reported in the original publication for (A) RNAP, (B) pol II, and (C) T7 pol [6,28,29]. Each coloured dot represents a single sample simulated from the posterior distribution of parameters/models for the respective polymerase. 30 samples were generated from each of the three posterior distributions. For RNAP, [NTP]_*eq*_ is defined as [ATP] = 5 *μ*M, [CTP] = 2.5 *μ*M, [GTP] = 10 *μ*M, and [UTP] = 10 *μ*M.

A large effective sample size (> 100 [54]) and a small 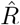 (< 1.1, as defined by Gelman et al. 1992 [55–57]) are essential for making reliable parameter estimates. Table 2 suggests that the parameters in the 95 % credible set of models are sufficiently estimated by these criteria.

These results indicate that the best models for the datasets examined are Models 11 and 12 for both RNAP and pol II, and Model 5 for T7 pol (Fig 4B).

For pol II, Model 12 has the highest posterior probability *P*(*M*_12_|*D*) = 0.71. This is the most complex model considered, with 6 estimated parameters. In Model 12 translocation, NTP binding and catalysis are all kinetic processes; the displacement required to facilitate formation of the translocation transition state, *δ*_1_ < *δ*, is estimated 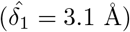; and the standard Gibbs energy of the posttranslocated state is influenced by parameter Δ*G*_*τ*1_ ≠ 0.

The posterior distribution for RNAP consists of the same set models as that of pol II. For RNAP, Model 11 has the highest probability *P*(*M*_11_|*D*) = 0.81. This model is a submodel of Model 12 with one fewer parameter: in Model 11 NTP binding is treated as an equilibrium process while in Model 12 it is not.

The only model in the 95 % credible set for T7 pol is Model 5 P(M_5_|D) = 0.96. In Model 5 (4 parameters) translocation, but not binding, is treated as an equilibrium process, and Δ*G*_*τ*1_ is estimated. This positions T7 pol in a quite different area of the model space to the other two polymerases.

### Translocation rates differ among RNA polymerases

For RNAP and pol II, we estimate that a partial equilibrium approximation for the translocation step is inadequate. The posterior probability that such models are inadequate is 1.00 (see Table 2). For T7 pol, however, translocation is significantly faster than catalysis and is best modelled with a partial equilibrium approximation.

Using estimates for 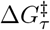 and Δ*G*_*τ*1_ under the maximum posterior models (Model 11 for RNAP and Model 12 for pol II) we estimate the mean forward 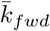 and backward 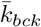 translocation rates averaged across the *rpoB* sequence as: 230 s^−1^ and 112 s^−1^ for RNAP, and 350 s^−1^ and 12.7 s^−1^ for pol II, respectively (3 sf). These estimates are within one order of magnitude of the respective estimate for the rate of catalysis (Fig 5) suggesting that translocation and catalysis indeed occur on similar timescales.

For RNAP and pol II, translocation has frequently been modelled as an equilibrium process [6,23,28,42,44], however in some recent analyses this assumption has been rejected [18,19,43,58,59]. Our Bayesian analysis supports this. In contrast, there is general agreement that translocation in T7 pol is adequately modelled as an equilibrium process [29,60,61].

### The data does not determine the kinetics of the NTP binding step

It remains unclear how to best model the NTP binding step. Models that describe NTP binding as a kinetic process have posterior probabilities of 0.19 for RNAP, 0.71 for pol II and 0.96 for T7 pol (Table 2). However, in our sensitivity analysis, where we used different a prior distribution for 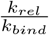, these probabilities were 0.65, 0.22, and 0.19, respectively (results not shown).

Furthermore, 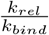 and *k_bind_* (Models 5 and 12) are unable to be estimated simultaneously. For pol II and for T7 pol, *k_bind_* is estimated at around 0.48 and 1.4 *μM*^−1^ s^−1^ respectively with fairly narrow 95% highest posterior density (HPD) intervals (Fig 5). However, the HPD interval of 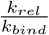 spans three orders of magnitude and the value of this parameter was therefore poorly informed by the data. For RNAP, in contrast, neither *k_bind_* nor 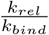 were well-informed by the data and both have HPD intervals spanning 1-2 orders of magnitude. This non-identifiability – where two or more parameters are unable to be estimated simultaneously (S4 Appendix) – highlights the appeal in an NTP binding equilibrium model where only one parameter 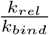 needs to be estimated, despite the unrealistic assumptions it may invoke. In the case of each enzyme, the data has taught us nothing about one or two of the binding parameters.

The pause-free mean velocities measured during transcription elongation follow Michaelis-Menten kinetics even though the reaction cycle is more complicated than that of a simple enzyme [62]. As such, the inability to resolve the timescale of the substrate binding step is unsurprising [63–65].

Overall, these three factors, i) the sensitivity of the posterior probabilities to the choice of prior, ii) the intermediate magnitude of said probabilities, iii) the inability to estimate both 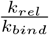 and *k_bind_* simultaneously, collectively imply the data carries very little information about the rates of NTP binding and release.

In the transcription literature, NTP binding is almost always assumed to achieve equilibrium for RNAP, pol II, and T7 pol [6,18,19,23,28,29,39,42,43,61]. However Mejia et al. 2015 [44] have shown that NTP binding is indeed rate-limiting, and that mutations in the RNAP trigger loop impair the binding rate thus suggesting that the trigger loop is coupled with NTP binding.

### RNAP has an energetic preference for the posttranslocated state

In previous stochastic sequence-dependent models [18,23] the standard Gibbs energies of the pre and posttranslocated states have been based solely on the nucleic acid basepairing energies. Our models include an additional term, Δ*G*_*τ*1_, to account for potential interactions between the protein and the nucleic acid. The marginal posterior probability of a model in which an additional term Δ*G*_*τ*1_ is required is 1.00 in all three polymerases. In each case Δ*G*_*τ*1_ was estimated to be less than 0 *k_B_T* and 0 *k_B_T* is not included in the 95 % HPD interval (Fig 5). We find that 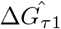 is the most significant in pol II and T7 pol: −4.6 *k_B_T* and −4.0 *k_B_T* respectively, while 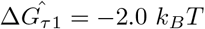 for RNAP (2 sf).

These results suggest that structural elements within RNA polymerases can energetically favour posttranslocated states over pretranslocated states. We note that the sequence-dependent contribution of the dangling end of the DNA/RNA hybrid is included in the thermodynamic model. The energetic bias for the posttranslocated state is separable from this effect.

To facilitate comparison with previous deterministic models, using our estimates of Δ*G*_*τ*1_ we calculated the equilibrium constant between the pre and posttranslocated states. Geometrically averaged across the *rpoB* gene, these are

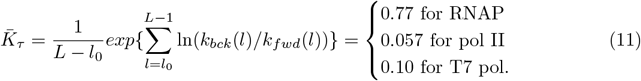

Thus, for all three polymerases, *K_τ_* < 1, indicating that the small energetic preference that the protein has for the posttranslocated state is sufficient to override the loss of basepairing energy, thereby biasing the system towards population of the posttranslocated positions. This is in agreement with estimates made for pol II and T7 pol [28,29,37,38,42] and Kireeva et al. 2018 [59] for RNAP: *“forward translocation occurs in milliseconds and is poorly reversible”*. However these estimates are inconsistent with some RNAP and pol II studies which place this ratio above 1 [6,19,43,53].

Kinetic modelling can itself suggest no physical mechanism for the stabilisation. Yu et al. 2012 [38] have identified a conserved tyrosine residue near the active site of T7 pol that pushes against the 3′ end of the mRNA, and thus stabilises the posttranslocated state. They propose a similar mechanism for the multi-subunit RNA polymerases.

### *δ*_1_ may be an important parameter but its physical meaning is unclear

Our results suggest that *δ*_1_, the distance that RNA polymerase must translocate forward by to reach the translocation transition state, is a necessary parameter to estimate for RNAP and pol II. Setting *δ*_1_ = *δ*/2 is not sufficient. The marginal posterior probability of models which estimate this term is 1.00. *δ*_1_ is irrelevant to the modeling of the T7 pol data because the best models invoke a partial equilibrium approximation for the translocation step.

While our prior distribution restricted *δ*_1_ to lie in the range (0, *δ*), the upper end our 95% HPD intervals of *δ*_1_ for RNAP and pol II are very close to *δ* = 3.4 Å. If it was not for this prior distribution, *δ*_1_ estimates would have included values higher than *δ*. Similar results have been observed by Maoiléidigh et al. 2011 [19] for RNAP.

Our interpretation of *δ*_1_ implies it should never be greater than *δ* nor should *δ* be more than the width of one basepair. The physical meaning of *δ*_1_ with values greater than *δ* is thus unclear. It is noted that *δ*_1_ is only used when *F* ≠ 0.

### Comparing the kinetics of RNA polymerases

The *in vivo* rate of transcription elongation varies considerably across RNAP, pol II and T7 pol. The prokaryotic and eukaryotic RNA polymerases have a mean rate ranging from 20-120 bp/s [46,47,49,50,66–68], which may be slowed down in histone-wrapped regions of eukaryotic genomes [9]. In contrast, Bacteriophage T7 pol operates up to an order of magnitude faster (around 200-240 bp/s [50, 69]) and is known to be quite insensitive to transcriptional pause sites [11,29].

In additional to these differences, we have shown that translocation is very rapid in T7 pol, relative to the rate of NTP incorporation, while the disparity is much less significant in RNAP and pol II. Furthermore, the model does not fit the data for T7 pol as closely it does for RNAP and pol II (Fig 6). T7 pol therefore seems to operate under quite a different kinetic scheme than that of the cellular polymerases, which is not unexpected given their distant evolutionary relationship [5].

In general, the elongation velocity of RNA polymerase is significantly slower in an optical trap (with estimates ranging from 9.7-22 bp/s for RNAP [13–15,44,70]) compared with that of the untethered enzyme (with estimates *in vitro* or *in vivo* ranging from 25-118 bp/s for RNAP [46,50,71,72]). This relationship holds for multiple RNA polymerases including *E. coli* RNAP, *S. cerevisiae* pol II [42,43,53,73], Bacteriophage T7 pol [11,29,50,52], and Bacteriophage Φ6 P2 [12,74]. This suggests that optical trapping perturbs the system to a significant extent. Additionally, varying degrees of heterogeneity in elongation rate have been observed across different polymerase complexes even under the same conditions [13,15,29].

The velocity perturbations resulting from the optical trapping apparatus will be propagated into the model parameters, especially *k_cat_*, and 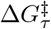, and some caution is needed when extrapolating these results to untethered systems.

### Bayesian inference of transcription elongation

To our knowledge we are the first to perform Bayesian inference on single-molecule models of transcription elongation. This was achieved by simulation which necessitated the use of approximate Bayesian computation. An alternative would be to build and use a likelihood function (ie. the probability of taking exactly *t* units of time for RNA polymerase to copy the sequence *n* times). The latter approach can be achieved using chemical master equations, as opposed to (Gillespie) sampling from the distribution. Finding analytical, stable numerical, or approximate solutions to the chemical master equations could provide a similar insight in less computational time, however is susceptible to a multitude of analytical and numerical issues associated with the exponentiation of an arbitrary transition rate matrix that grows with the length of the sequence (S2 Appendix) [75]. This problem would be amplified by the introduction of backtracking, hypertranslocation, or NTP misincorporation reactions into the model, for instance. The Bayesian framework we have presented, although computationally intensive due to its simulation requirement, is general and will work on any model of transcription without the need to resolve these issues. The path has been paved for modelling transcriptional pausing, for instance [18,23,76]. Nevertheless, likelihood-based Bayesian inference is an approach that should be explored in the future.

We have demonstrated that single-molecule data can be usefully analysed using a Bayesian inference and model selection framework. This analysis would have even greater statistical power if applied to the progression of individual RNA polymerase complexes instead of mean velocities averaged across multiple experiments.

## Conclusion

In this article we evaluated some simple Brownian ratchet models of transcription elongation (Fig 2). By varying the parameterisation of the translocation step (Fig 3) and incorporating partial equilibrium approximations commonly invoked in the literature (Fig 4A) we enumerated a total of 12 related models (Fig 4B). Using stochastic simulations and approximate Bayesian computation, we then assessed which of these models were capable of describing the force-velocity data previously measured for several RNA polymerases (Table 2 and Fig 5) using single-molecule optical trapping experiments [6,28,29].

Our analysis suggests that 1) different partial equilibrium approximations of the translocation step are appropriate for the multisubunit RNA polymerases versus the single subunit T7 RNA polymerase. 2) Treatment of the NTP binding step remains a point of ambiguity. The existing data does not place strong constraints on the modelling of this step. 3) There is an energetic bias for posttranslocated state. 4) The model of the force-dependent translocation, which invokes transition state theory, is not physically realistic.

## S1 Appendix. Stochastic simulation

Reactions are simulated using the Gillespie algorithm [26]. Given the current state *s* and a set of possible reactions *s* → *s*_1_, *s* → *s*_2_,…, *s* → *s_n_* with rate constants *k*_1_, *k*_2_,…, *k_n_*, the next reaction to perform is sampled proportional to its rate:

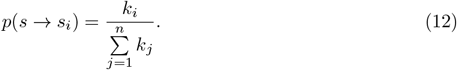

The amount of time the reaction takes to occur is sampled from the exponential distribution with rate 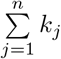.

## S2 Appendix. Chemical master equations

The chemical master equations for the four equilibrium variants (Fig 4A) of single nucleotide addition cycles are provided in this section. Transcription of a full gene involves chaining multiple of these single-cycle models together.

### Translocation and binding equilibrium model

The state pathway of a single-cycle of the translocation and binding equilibrium model is

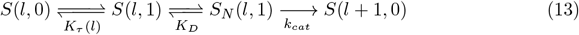

where *S_N_* denotes a state where NTP is bound. As states *S*(*l*, 0), *S*(*l*, 1), and *S_N_*(*l*, 1) are in mutual equilibrium they can be coalesced into one state. Let *S*(*l*) be the coalesced state that exists in equilibrium between the three, then the pathway may be rewritten as

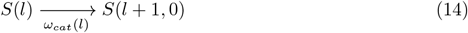

where *ω_cat_*(*l*) is the *effective* rate of catalysis from this state. This term is equal to *k_cat_* multiplied by the proportion of time that the coalesced state has NTP bound, ie. in state *S_N_*(*l*, 1).

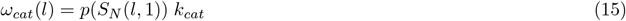

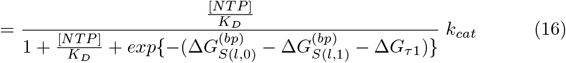

Let 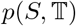 be the probability of the system existing in state *S* at time 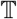. Under this model, the chemical master equation of a single-cycle is:

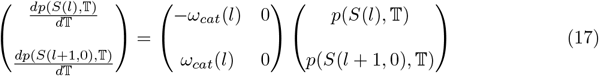

This system is simple enough to solve analytically. Let 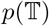 be the Markov transition matrix after time 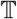 and let *Q* be the Markov process transition rate matrix. In these matrices, entry *i*, *j* is the probability (*p*) or rate (*Q*) of transition from *i* to *j*.

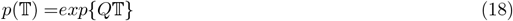

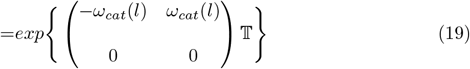

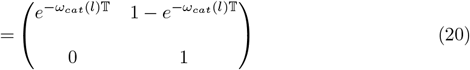

It is noted that as the time 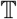 approaches infinity, the probability of the system existing in state *S*(*l* + 1, 0) approaches 1, because it is an absorbing state.

Let 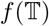 be the probability density of taking *exactly* 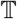 units of time to arrive at state *S*(*l* + 1, 0), starting from *S*(*l*). Because *S*(*l* + 1,0) is an absorbing state, computing 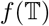 is trivial.

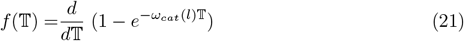

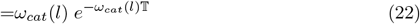

Transcribing the full gene requires the traversal of all *L* – *l*_0_ + 1 states. The size of the transition rate matrix for transcribing the entire sequence therefore grows with the length of the DNA sequence.

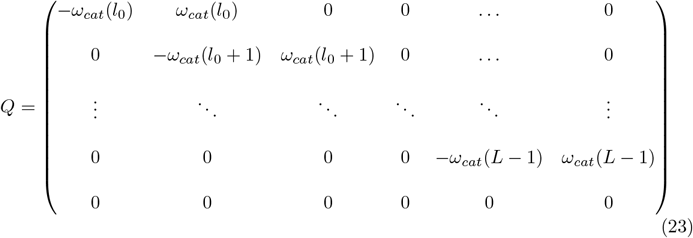

Applying the matrix exponential function to a linear pathway, such as the one described above, has an analytical solution [77]. However, for non-linear systems with an arbitrary number of states (eg. the other 3 equilibrium model variants), analytical solutions may not exist and numerical solutions likely exhibit numerical instabilities [75]. This option was not further investigated in this project (simulation was used instead). However, analytical or stable numerical solutions for 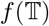 for an arbitrary model would facilitate the use of likelihood functions in Bayesian inference, thereby rendering simulation obsolete.

### Binding equilibrium model

The state pathway of a single-cycle of the binding equilibrium model is

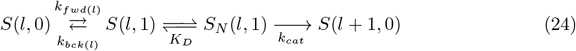

The binding equilibrium assumption permits the coalescence of *S*(*l*, 1) and *S_N_*(*l*, 1) into a single state *S_N_*(*l*). Thus, the pathway can be rewritten as

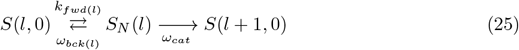

where *ω*_*bck*(*l*)_ is the effective rate of backwards translocation from *S_N_*(*l*), and *ω_cat_* is the effective rate of catalysis. These rates are derived by multiplying their composite rates – *k*_*bck*(*l*)_ and *k_cat_* – by the probability of the system existing in the required state to apply the reaction – *p*(*S*(*l*, 1)) and *p*(*S_N_*(*l*, 1)).

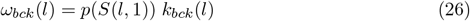

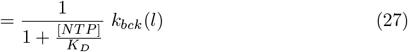

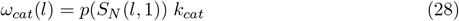

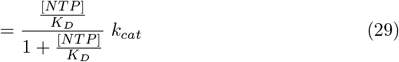

Let 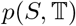 be the probability of the system being at state *S* at time 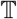. Under this model, the chemical master equation of a single-cycle is:

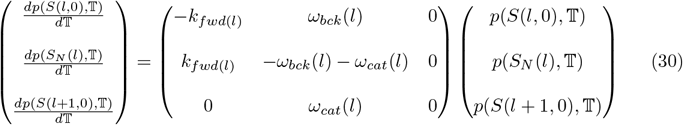

### Translocation equilibrium model

The state pathway of a single-cycle of the translocation equilibrium model is

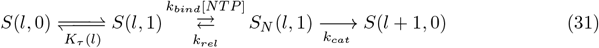

The translocation equilibrium assumption permits the coalescence of *S*(*l*, 0) and *S*(*l*, 1) into a single state *S_τ_*(*l*). Thus, the pathway can be rewritten as

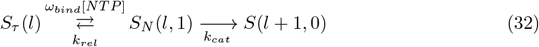

where *ω_bind_* is the effective rate of NTP binding.

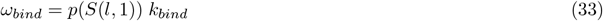

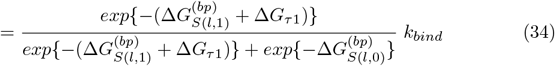

Let 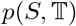 be the probability of the system being at state *S* at time 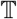. Under this model, the chemical master equation of a single-cycle is:

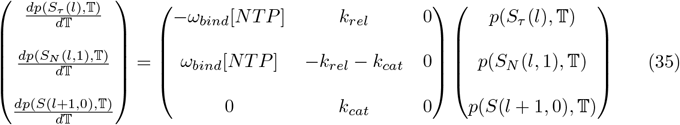

### Full kinetic model

The state pathway of a single-cycle of the full kinetic model is

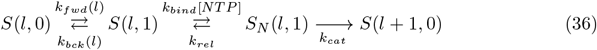

Let 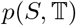 be the probability of the system being at state *S* at time 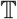. Under this model, the chemical master equation of a single-cycle is:

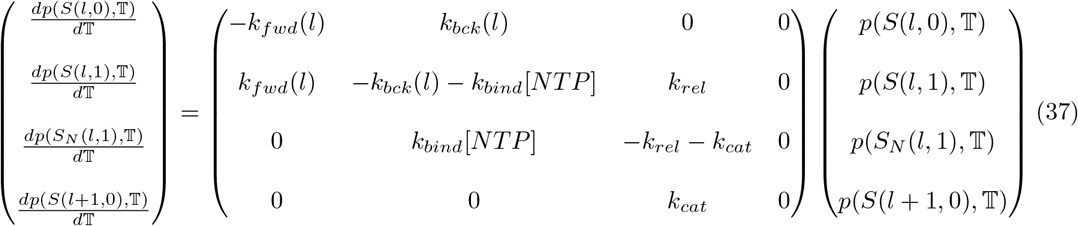

## S3 Appendix. MCMC-ABC

Given model/parameters Θ and observed data *D* = (*D*_1_, *D*_2_,…, *D_n_*), Bayesian inference conventionally involves approximating the posterior probability distribution *P*(Θ|*D*) using the likelihood *P*(*D*|Θ) and the prior *P*(Θ).

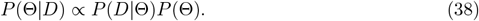

As there is no easily computed likelihood function, simulation is used. The chi-squared test statistic *X*^2^ evaluates how well a given set of parameters fits the data.

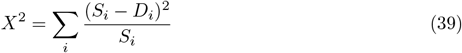

where *S_i_* is the mean velocity simulated under the same [NTP] and applied force *F* that *D_i_* was measured under. The probability that *X*^2^ = 0 is equal to the likelihood *P*(*D*|Θ), however it is computationally impractical to only accept parameters into the posterior distribution when the simulation yields *X*^2^ = 0. Therefore a threshold *ϵ* is used and sample Θ_*i*_ is accepted into the posterior only if *X*^2^(Θ_*i*_) ≤ *ϵ*. This method is called approximate Bayesian computation [27,45]. This is coupled with Markov chain Monte Carlo (MCMC) to give the MCMC-ABC algorithm which is becoming increasingly popular among computational biologists [27,78].

Each MCMC chain estimated six parameters and the model indicator *M*. This means the 12 models share the same parameter objects in the MCMC. There are therefore seven terms to estimate: *M*, *k_cat_*, 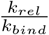, *k_bind_*, Δ*G*_*τ*1_, 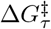, and *δ*_1_.

When the current model *M_i_* does not use a parameter (eg. in Model 5 *δ*1 is not used), the parameter is still estimated even though it is not being used. When the model requires a parameter to be held constant (eg. in Model 1 Δ*G*_*τ*1_ = 0), the parameter is set to its constant during the simulation. This is done without affecting or being affected by its current estimate, which is used by other models.

To achieve convergence, we used an exponential cooling scheme on *ϵ* [78] where *ϵ*_*i*+1_ = max(*ϵ_min_*, *ϵ_iγ_*) for manually tuned values of 0 < *γ* < 1 and *ϵ*_0_. Chains which failed to converge were discarded. A heavy-tailed distribution [79] is used as a proposal function, and the parameter to change at each step in the MCMC is selected uniformly at random.

We ran one or more independent MCMC-ABC chains for each selected *ϵ_min_* / RNA polymerase combination. Selecting the threshold *ϵ_min_* is a critical process in approximate Bayesian computation. Threshold *ϵ_min_* must be large enough to achieve convergence within finite computational resources, but small enough that the resulting posterior distribution is still an accurate approximation of the true posterior distribution. For each RNA polymerase we set *ϵ_min_* to some initial guess. Then we ran the MCMC chain until the ESS for *X*^2^ was large (> 300) and lowered *ϵ_min_* to the bottom 0.05 quantile of the posterior distribution of *X*^2^. This step was repeated until either: a) the distribution of model indicators *M* converged (model posterior probabilities have changed by less than 0.01, on average). Or, b) the acceptance rate was less than 5%. The values of *ϵ_min_* used in the final posterior distributions were 2.39 for RNAP, 0.705 for pol II, and 4.63 for T7 pol (Table 2).

Parameter point estimates (Fig 5) are the geometric median: that is the value which minimises the total Euclidean distance from the other posterior samples. Parameters were normalised into z-scores first. Our code is open source and available at http://www.polymerase.nz. Textfiles containing the posterior distributions and simulation settings are available to download or visualise with the software.

## S4 Appendix. Prior distributions

### Prior for 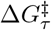, which governs the rates of translocation

RNAP/pol II: to select a prior for 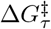 we simulated transcription on the *rpoB* gene under Model 3 – the simplest binding equilibrium model. 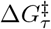 and *k_cat_* were sampled uniformly from a relevant range, with *K_D_* held constant at 100 *μ*M and [NTP] = 1000 *μ*M. For each simulation, the mean elongation velocity was calculated. The results are displayed in S1 Fig.

This plot shows that as the energy barrier of translocation 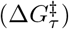 increases, the velocity decreases. If 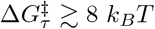 then it becomes impossible to achieve a realistic mean velocity, providing a relatively clear upper bound on this parameter. If 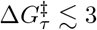 *k_B_ T* then translocation becomes very rapid and the same distribution of velocities is obtained in simulations, irrespective of the exact value of 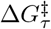. In this case catalysis becomes strongly rate-limiting, and it would be appropriate to apply a partial equilibrium approximation to the translocation step. This provides an effective lower bound for parameter 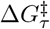. Therefore we centered our prior distribution for 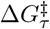 in this interval (a normal distribution with a mean of 5.5 and a standard deviation of 0.97, so that the central 99% interval is (3, 8)). We performed the same analysis with different values of *K_D_*, as well as varying Δ*G*_*τ*1_, and arrived at the same interval for 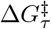 (results not shown).

T7 pol: the same analysis was performed, however with 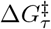 at its prior mean of −3.3 *k_B_T* (S1 Fig).

### Prior for *k_bind_*, which governs the rate of NTP binding

To select a prior for *k_bind_* we performed similar simulations, but instead used Model 2 – the simplest kinetic binding model. *k_bind_* and *k_cat_* were sampled uniformly from relevant ranges, *K_D_* was set to 100 *μ*M and [NTP] = 1000 *μM*. (S1 Fig).

Depending on the exact value of *k_cat_*, if *k_bind_* ≲ 0.1 *μ*M^−1^ s^−1^, then it is impossible to achieve a realistic velocity, providing a relatively clear lower bound on this parameter. If *k_bind_* ≳ 5 *μ*M^−1^ s^−1^ then binding becomes very rapid and the same distribution of velocities is obtained in simulations, irrespective of the exact value of *k_bind_*. Again this is because catalysis becomes strongly rate limiting in this region, and it would be appropriate to apply a partial equilibrium approximation to the binding step. Hence we centered our (lognormal) prior around the interval (0.01, 5) – the conservatively selected bounds reflecting that the experimental data has been collected at differing NTP concentrations, altering the rate. Performing the same analysis with different parameters gave us a similar prior.

### Prior distribution related to rate of NTP release

A model is non-identifiable if two or more parameterisations can produce the same output. Our preliminary results suggested non-identifiability between 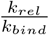 and *k_bind_* (S1 Fig). When *k_bind_* is low (and hence binding is rate-limiting), there is an approximately linear relationship between 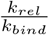 and *k_bind_*. As *k_bind_* increases from 0, the dissociation constant 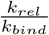 must also increase in order for the system to achieve the same velocity. However, as binding comes closer to achieving equilibrium, 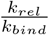 converges. Most previous estimates of *K_D_* have assumed binding to be at equilibrium. This assumption restrains the values which *K_D_* may take, and subsequently estimates for *K_D_* are typically in the order of 10^1^ – 10^2^ *μ*M. However for a model in which binding is slow it is expected that estimates of 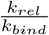 can be lower. This has indeed been demonstrated by Mejia et al. 2015 [44] who estimated 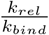 to be 0.6 *μ*M. Therefore the prior distribution for 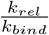 must permit both of these binding models to be tested fairly during Bayesian inference. We centered our lognormal prior for 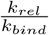 around a very broad range, with a central 95% interval of (0.2, 200).

It is noted that selecting a prior distribution which does not discriminate between the kinetic and equilibrium binding models *a priori* may not be plausible.

**S1 Fig. Simulations of the elongation pathway**. Each point is a single simulation of the full *rpoB* gene (4029 nt). For (A-C), Parameters on the x- and z-axis are sampled uniformly at random from the displayed range at the beginning of each trial. The y-axis of each plot (mean elongation velocity) is then measured from the respective simulation. [NTP] and *F* held constant at 1000 *μ*M and 0 pN respectively. (A) and (B): Relationship between 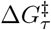 and *k_cat_* for the melting model with binding at equilibrium (Model 8). Δ*G*_*τ*1_ set to its prior mean (0 for RNAP and pol II, and −3.3 for T7 pol). (C) Relationship between *k_bind_* and *k_cat_* for the kinetic binding model with translocation at equilibrium (Model 2). (D) Relationship between *K_D_* and *k_bind_* with translocation held at equilibrium (Model 2). *K_D_* and *k_bind_* sampled uniformly from specified range and velocity is measured. Samples with simulated velocities outside of the range 1-2 bp/s were discarded. [NTP] = 10 *μ*M and *k_cat_* = 100 s^−1^.

## Acknowledgments

We wish to acknowledge the contribution of NeSI high-performance computing facilities to the results of this research. NZ’s national facilities are provided by the NZ eScience Infrastructure and funded jointly by NeSI’s collaborator institutions and through the Ministry of Business, Innovation & Employment’s Research Infrastructure programme. URL https://www.nesi.org.nz.

