## Supplementary figures and images for "Bayesian inference and comparison of stochastic transcription elongation models"

### S1 Fig

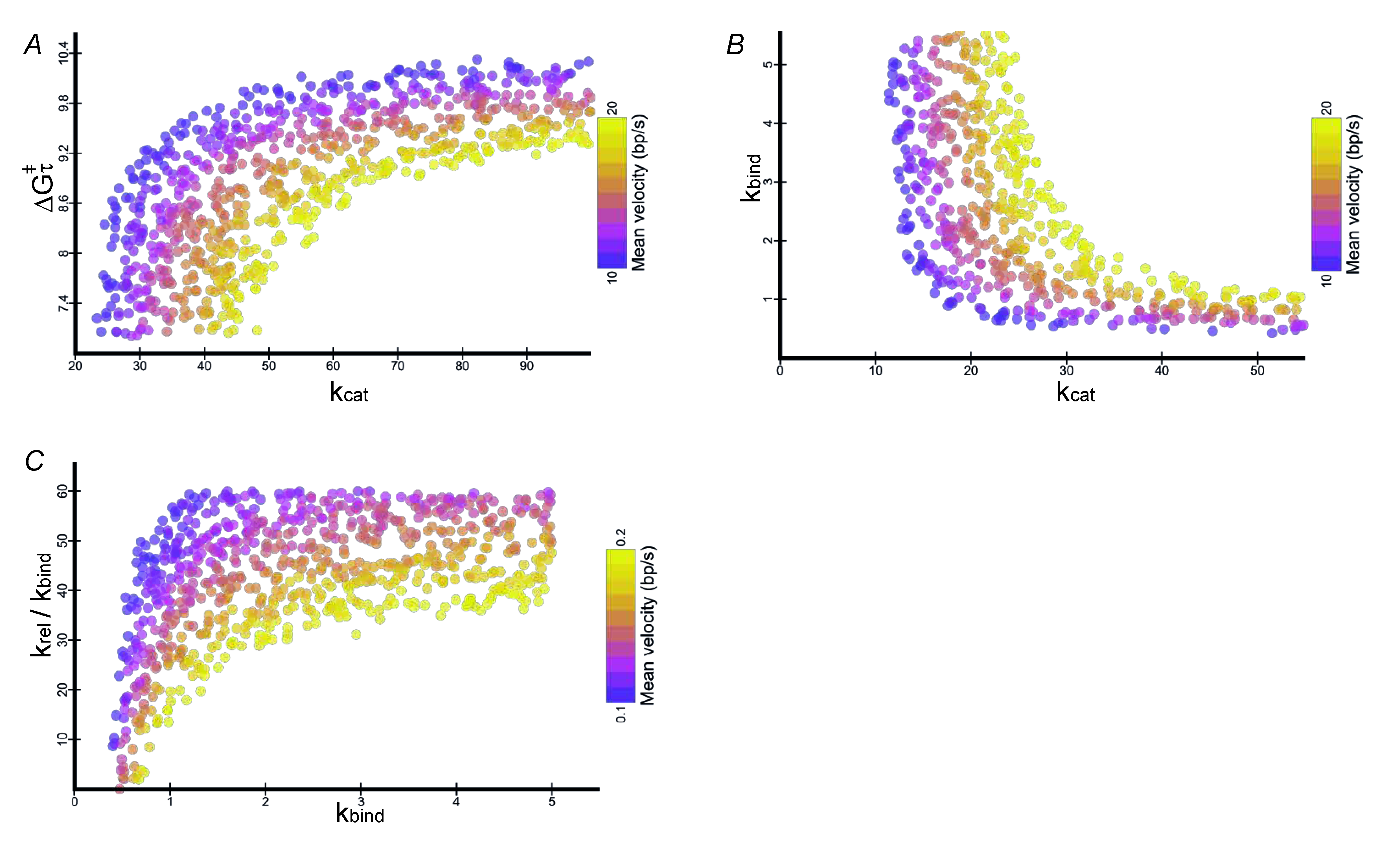
